# Cinematic Simulation of Substrate-to-Product Chemical Reactions with Chemical Accuracy Using QuantaMind MD

**DOI:** 10.1101/2025.09.28.679094

**Authors:** Song Xia, Deqiang Zhang

## Abstract

Polyethylene terephthalate hydrolases (PETases) are enzymes that catalyze the breakdown of PET plastic. Previous studies have employed classical molecular dynamics (MD) and quantum mechanical/molecular mechanical (QM/MM) simulations to probe the catalytic mechanisms of the PETase-catalyzed reaction. In this work, we apply QuantaMind, a deep learning-based machine learning force field (MLFF) trained on diverse chemical systems and transition-state energies at density functional theory (DFT)-level, to simulate the complete catalytic cycle of the PETase reaction. Our simulations capture key proton transfer events and the stabilization role of the oxyanion hole in PETase, in agreement with previous computational studies, without introducing artificial biases to the interactions. The simulations also provide estimates of free energy barriers for each reaction step with uncertainty estimation, consistent with experimental values and previous QM/MM studies. To our knowledge, this work represents the first complete DFT-level MD simulation of a biomolecular enzymatic reaction using DL-based MLFFs.

## Introduction

Polyethylene terephthalate (PET) is the most common thermoplastic polymer of the polyester family, widely used in textiles and food and beverage containers.^1^ As a result, vast quantities of PET waste accumulate in the environment, posing a severe threat to global ecosystems. In 2016, a PET-assimilating bacterium, *Ideonella sakaiensis* 201-F6, was isolated from a PET recycling facility. This bacterium secretes a cutinase-like enzyme, IsPETase, that degrades PET to mono (2-hydroxyethyl) terephthalic acid (MHET), terephthalate (TPA), and ethylene glycol at ambient temperature.^2^ Since then, efforts have been devoted to engineering this enzyme, resulting in numerous improved variants.^3–13^

The catalytic mechanisms of serine hydrolases, including PETase, have been extensively studied. Conflicting hypotheses have been proposed with particular disagreement regarding proton transfer events during the catalytic cycle.^14-18^While QM/MM studies have provided valuable insights, these methods have inherent limitations, such as boundary issues and limited accuracy in the MM component, and therefore introduce artifacts and produce different results.^19–26^ Additionally, QM/MM simulations often require a large QM region for converged results, which is computationally expensive.^27^

Machine learning force fields (MLFFs) provide a promising alternative by mapping chemical descriptors to the quantum mechanical energy of systems,^28^ combining the efficiency of classical force fields with the accuracy of QM methods. As deep learning (DL)^29^ transforms computational chemistry, several DL-based MLFFs have been developed for MD simulations.^30–33^ Previously, we developed QuantaMind,^34^ a DL-based MLFF trained on diverse biomolecular fragments with an emphasis on transition-state geometries, shown to be effective in simulating reactions in aqueous solutions.

In this work, we demonstrate the application of QuantaMind in simulating the complete catalytic cycle of PETase. Our simulations elucidate key proton transfer events that are consistent with previous QM/MM studies. We estimate free energy barriers of 22 and 20 kcal/mol for the acylation and deacylation steps, respectively, along the lower bound path.

These values are close to the estimated experimental value of 18.7 kcal/mol^2,24^ and the calculated value of 21.1 kcal/mol by Boneta et al.^24^ To our knowledge, this research represents the first complete DFT-level MD simulation of a biomolecular enzymatic reaction enabled by a DL-based MLFF.

## Results and discussion

Figure 1 provides an overview of the enzymatic reaction simulated in this study. The simulation box (Figure 1A) contains 17,792 atoms in total, comprising the PETase protein (257 residues), one MHET substrate molecule, and 4,898 water molecules. Figure 1B illustrates the chemical reaction simulated: the hydrolysis of MHET to produce terephthalic acid and ethylene glycol.

**Figure 1:**
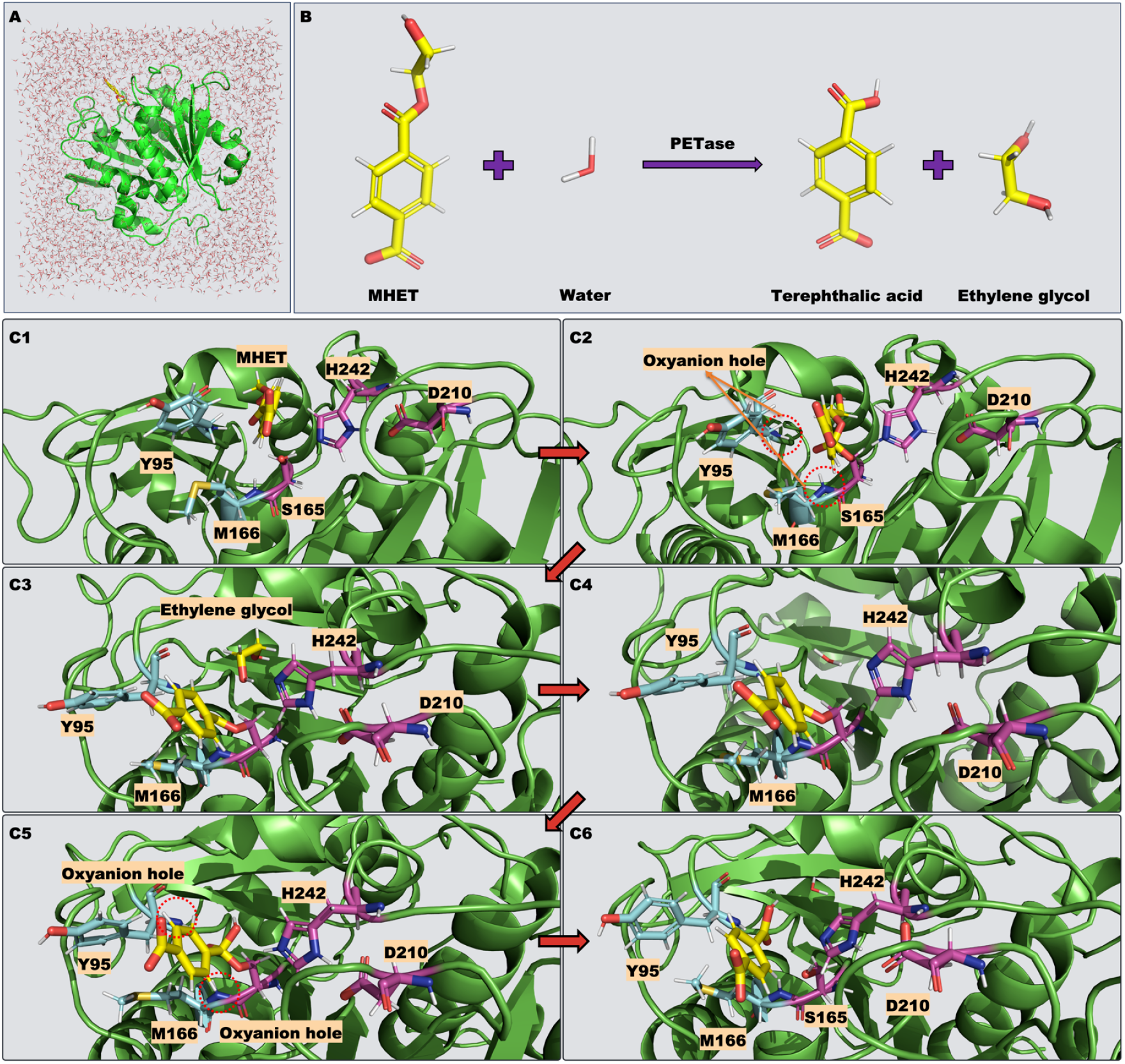
Overview of the PETase catalytic cycle simulated in this work. (**A**.) The simulation box used in this work. (**B**.) The chemical reaction studied in this work. (**C1-6**.) Representative frames from the simulation, indicating significant intermediates and products. For clarity, only the water molecule directly involved in the reaction is shown, while other solvent molecules are hidden.

Figures 1C1-6 present representative frames from the simulation trajectories, highlighting key intermediates along the catalytic pathway. Figure 1C1 shows in the initial state the substrate MHET positioned within the enzyme’s active site. After a nucleophilic attack by residue S165, a covalent intermediate forms between S165 and the reactant, as shown in Figure 1C2. Subsequent cleavage of the carbon-oxygen bond yields one of the reaction products, ethylene glycol and the acyl-enzyme intermediate (AEI) (Figure 1C3). As ethylene glycol departs, a water molecule enters the catalytic center (Figure 1C4) and performs a nucleophilic attack, forming a second tetrahedral intermediate (Figure 1C5). Finally, cleavage of the carbon-S165 oxygen bond results in the formation of terephthalic acid, restoring the catalytic triad to its original protonation state (Figure 1C6). Detailed analyses of the simulation trajectories are presented in the following subsections.

## PETase Acylation Reaction

During the acylation step of the PETase catalytic cycle, the serine residue S165 performs a nucleophilic attack on the ester bond of the MHET substrate, leading to its cleavage and the formation of ethylene glycol and the AEI. This acylation process proceeds through two distinct steps, depicted in Figures 2 and 3.

**Figure 2:**
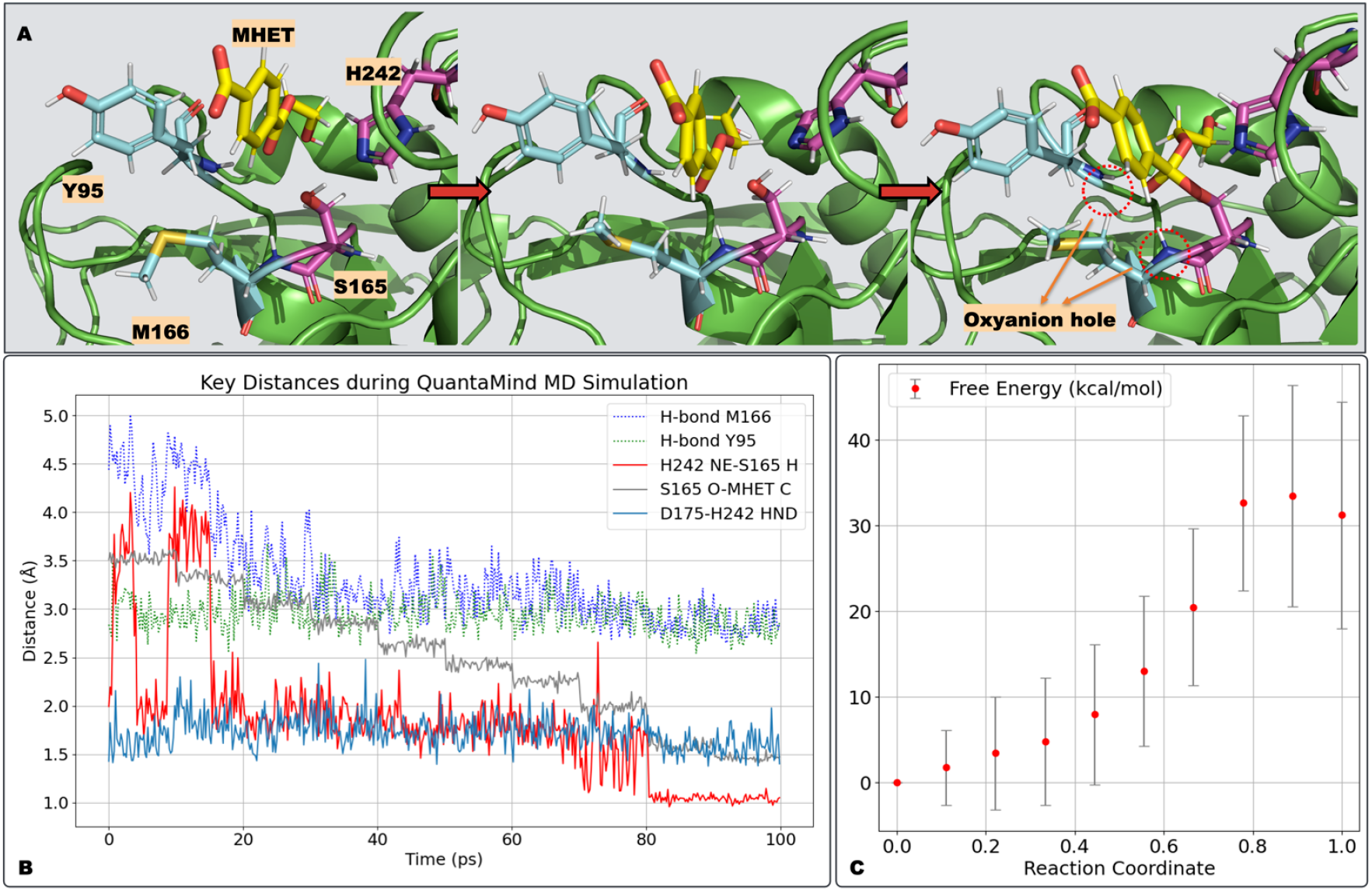
Simulation of the nucleophilic attack during PETase acylation reaction. (**A**) Visualization of the nucleophilic attack by the oxygen atom in S165 on the carbonyl carbon.(**B**) Key distances during the simulation. H-bond M166 and H-bond Y95 are the hydrogen bond distances between the carbonyl oxygen and the backbone nitrogen atoms of M166 and Y95, respectively. H242 NE-S165 H represents the distance between the N_*ϵ*_ of H242 and the S165 proton. S165 O-MHET C is the distance between the S165 oxygen and the carbonyl carbon in MHET. D175-H242 HND is the minimum distance between the two oxygen atoms in D175 and the hydrogen atom on N_*δ*_ of H242. (**C**) Free energy difference during the reaction, with red dots showing the average free energy difference relative to the initial state and error bars representing estimated standard deviations.

**Figure 3:**
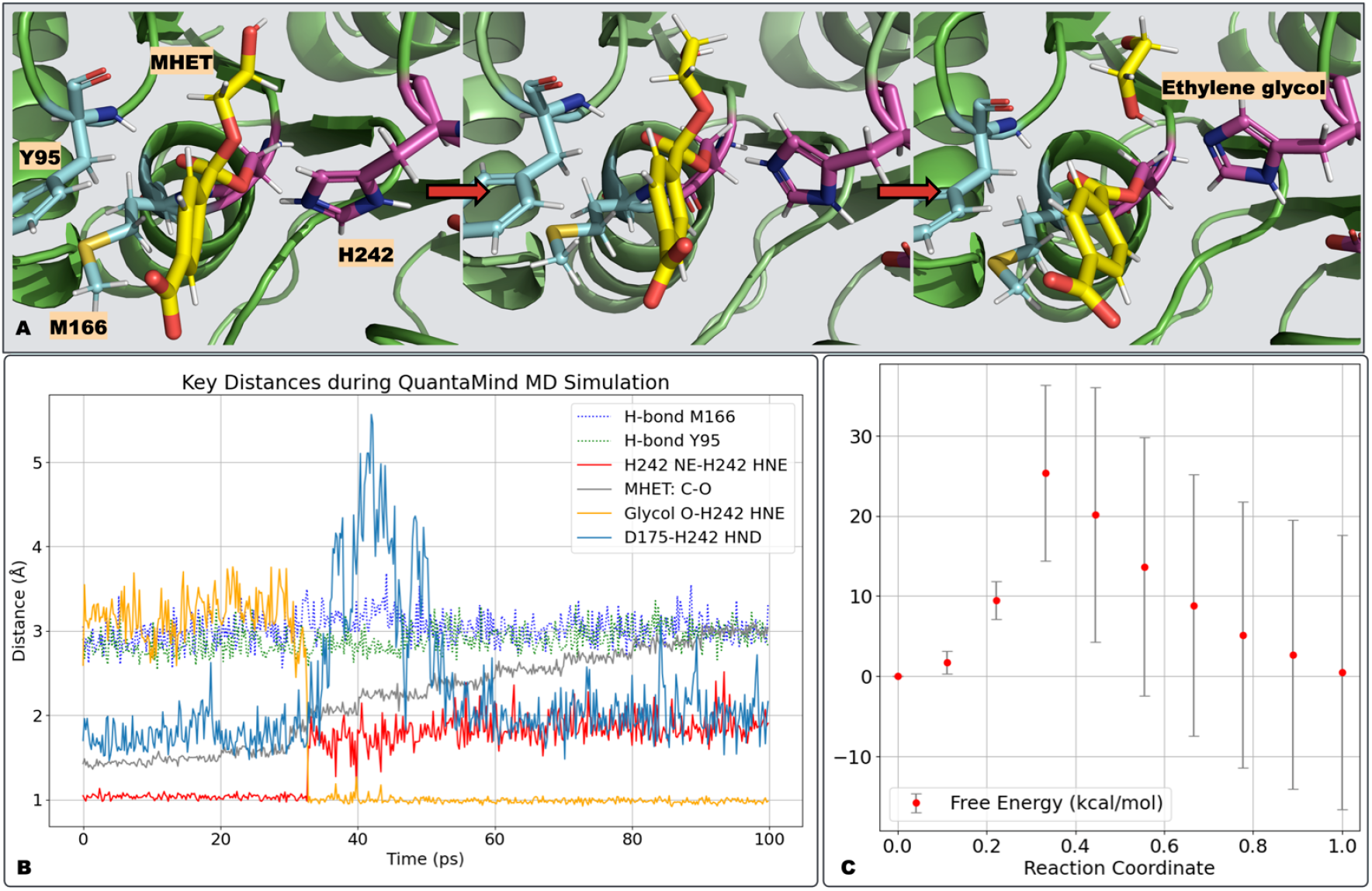
Simulation of the C-O bond cleavage during the PETase acylation reaction. (**A**) Visualization of the ester bond cleavage, leading to the formation of ethylene glycol and the AEI. (**B**) Key distances during the simulation. H-bond M166 and H-bond Y95 represent the hydrogen bond distances between the carbonyl oxygen and the backbone atoms of M166 and Y95, respectively. H242 NE-H242 HNE denotes the distance between the N_*ϵ*_ of H242 and the proton transferred from S165 in the previous step (Figure 2). MHET C-Water O is the ester bond distance in MHET. Glycol O-H242 HNE indicates the distance between H242’s proton and the glycol oxygen. D175-H242 HND is the minimum distance between the two oxygen atoms in D175 and the hydrogen atom on N_*δ*_ of H242. (**C**) Free energy difference during the reaction, with red dots showing the average free energy difference relative to the initial state and error bars representing estimated standard deviations.

Figure 2A presents representative snapshots from the nucleophilic attack simulation. Key interactions during this 100 ps simulation are quantified by monitoring several relevant distances (Figure 2B). The hydrogen bonds labeled as H-bond M166 and H-bond Y95 refer to the distances between the carbonyl oxygen and the backbone amide nitrogens of M166 and Y95, respectively. These atoms form the “oxyanion hole,” which stabilizes the tetrahedral intermediate during the reaction. Early in the trajectory, when S165 is distant from the MHET substrate, these hydrogen bond distances fluctuate substantially, particularly for H-bond M166. As the reaction proceeds, both distances stabilize around 3 Å, indicating stronger hydrogen bonds.

Another critical event is the proton transfer from S165 to H242 (denoted as H242 NE-S165 H). A proton jump is noted at approximately 80 ps. Prior to the jump, the distance between the N_*ϵ*_ of H242 and the S165 proton is around 2 Å, typical for a non-covalent hydrogen bond. At approximately 80 ps, this distance decreases and stabilizes around 1 Å, indicating the formation of a covalent N-H single bond.

QuantaMind MD also provides free energy barriers during the simulation, along with uncertainty estimates. As shown in Figure 2C, the red dots represent the average free energy difference relative to the initial state, with error bars indicating the estimated standard deviations. The free energy barrier for this step ranges from 22 kcal/mol (lower bound path) to 47 kcal/mol (upper bound path).

It is important to note that only the S165 O-MHET C distance–the distance between the S165 oxygen and the carbonyl carbon in MHET–is constrained using our enhanced sampling technique (Section QuantaMind MD Simulation), with no additional biases applied during the simulation.

In Figure 3A, representative snapshots from the ester bond cleavage simulation are shown. Key interactions are similarly analyzed by plotting relevant distances over this 100 ps simulation (Figure 3B). The hydrogen bonds, H-bond M166 and H-bond Y95, function as the “oxyanion hole,” stabilizing the tetrahedral intermediate and the AEI carbonyl oxygen.

A critical event during this simulation is the proton transfer from H242 to the nascent ethylene glycol product. As the ester bond breaks, the oxygen atom of the partially formed ethylene glycol requires an additional proton to become stable, capturing the proton from HIP242 (originating from S165) to form stable ethylene glycol. The Glycol O-H242 HNE and H242 NE-H242 HNE distance plots in Figure 3B illustrate this event: at around 35 ps, the proton migrates from H242 to the ethylene glycol oxygen, stabilizing at around 1 Å, indicative of a covalent O-H bond. Concurrently, the original covalent O-H bond elongates to approximately 1.8 Å, consistent with the non-covalent hydrogen bond distance.

Figure 3C shows free energy changes and uncertainty estimation during the simulation. The red dots represent the average free energy difference relative to the initial state, with error bars indicating the estimated standard deviations. The free energy barrier for this step ranges from 14 kcal/mol (lower bound path) to 37 kcal/mol (upper bound path).

During the enhanced sampling, only the MHET: C-O distance–the ester bond within the MHET molecule–is constrained (Section QuantaMind MD Simulation), with no additional biases applied.

## PETase Deacylation Reaction

During the deacylation reaction of PETase, a water molecule attacks the ester bond of the AEI, followed by cleavage of the ester bond between the reactant and S165. This regenerates the catalytic triad and releases terephthalic acid. The process proceeds through two steps, illustrated in Figures 4 and 5.

**Figure 4:**
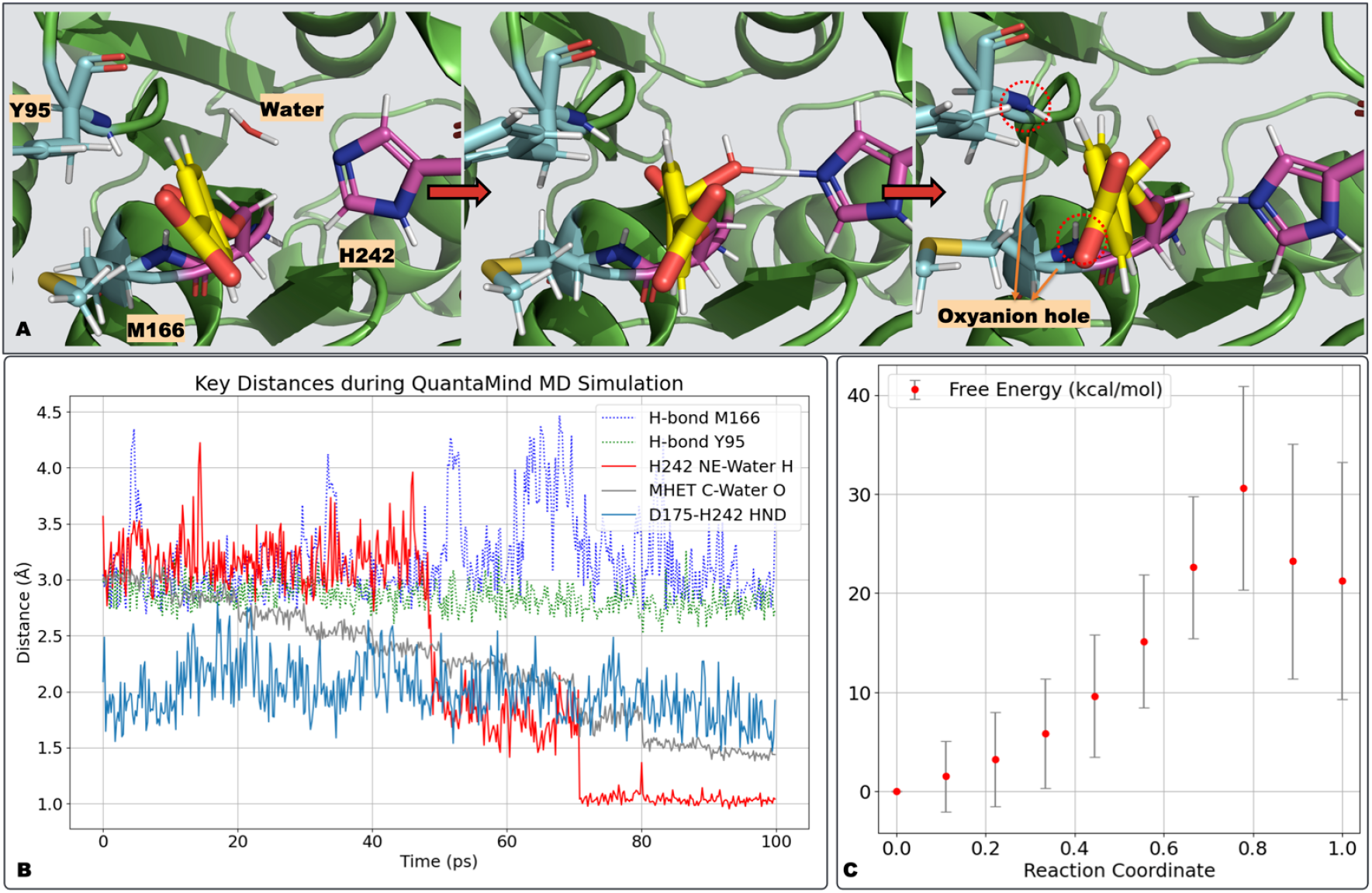
Simulation of the nucleophilic attack by a water molecule during the PETase deacylation reaction. (**A**) Visualization of the nucleophilic attack by a water molecule on the carbonyl carbon. (**B**) Key distances during the simulation. H-bond M166 and H-bond Y95 denote the hydrogen bond distances between the carbonyl oxygen and the backbones of M166 and Y95, respectively. H242 NE-Water H is the distance between the N_*ϵ*_ of H242 and the proton of water. MHET C-Water O is the distance between the water oxygen and the carbonyl carbon in the covalent intermediate. D175-H242 HND is the minimum distance between the two oxygen atoms in D175 and the hydrogen atom on N_*δ*_ of H242. (**C**) Free energy difference during the reaction, with red dots showing the average free energy difference relative to the initial state and error bars representing estimated standard deviations.

**Figure 5:**
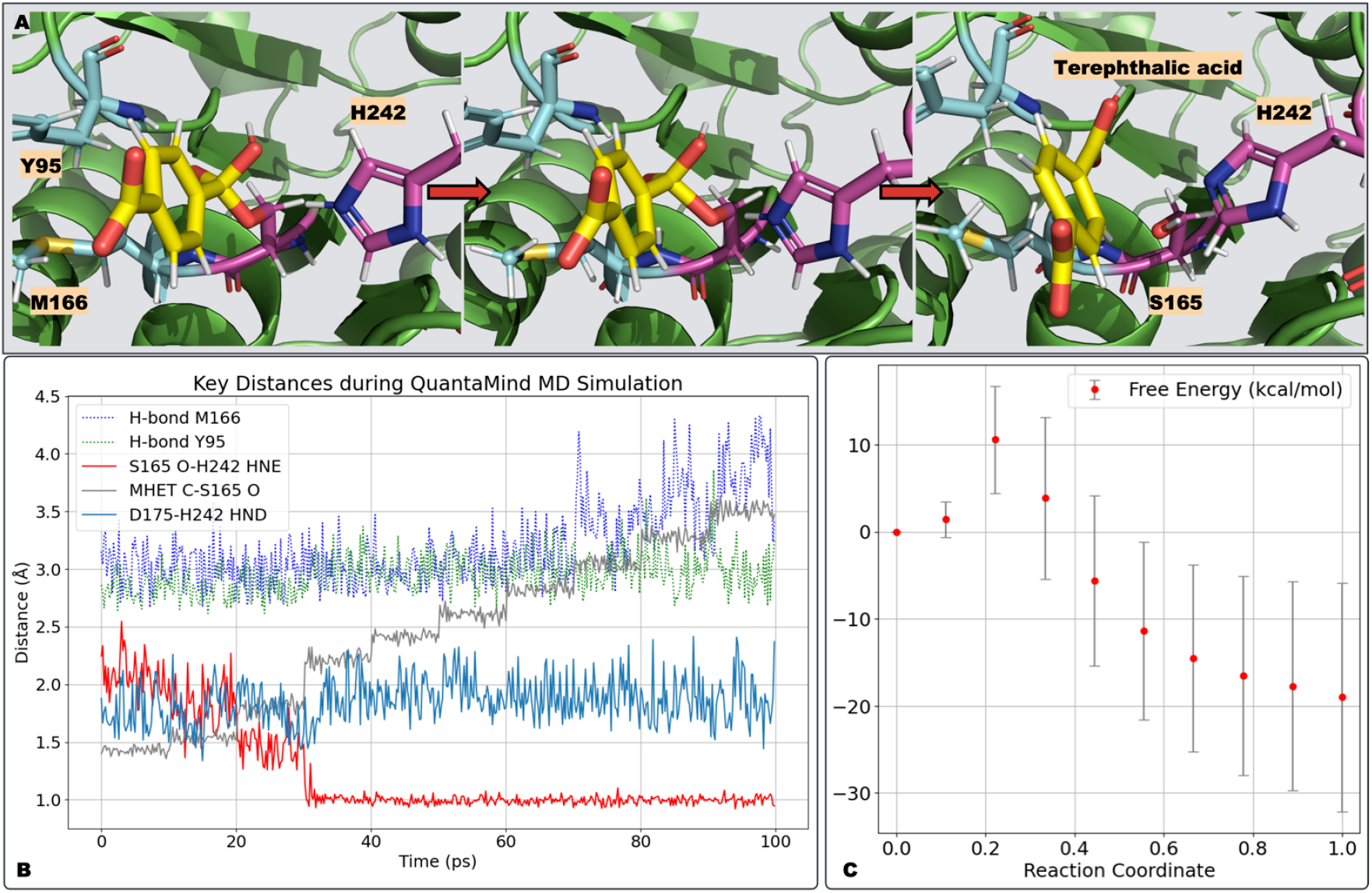
Simulation of the C-O bond cleavage during the PETase deacylation reaction.(A)Visualization of the ester bond cleavage, forming terephthalic acid and restoring the original catalytic triad. (**B**) Key distances during the simulation. H-bond M166 and H-bond Y95 represent the hydrogen bond distances between the carbonyl oxygen and the backbones of M166 and Y95, respectively. S165-Proton is the distance between the oxygen of S165 and the proton transferred from H242 in the previous step (Figure 4). MHET C-S165 O is the distance of the ester bond in the covalent intermediate. D175-H242 HND is the minimum distance between the two oxygen atoms in D175 and the hydrogen atom on N_*δ*_ of H242. (**C**) Free energy difference during the reaction, with red dots showing the average free energy difference relative to the initial state and error bars representing estimated standard deviations.

Figure 4A shows representative snapshots from the simulation of the water molecule’s nucleophilic attack. Key interactions are examined by plotting distances over this 100 ps simulation (Figure 4B). H-bond M166 and H-bond Y95 denote the hydrogen bond distances from the carbonyl oxygen to the backbone amide nitrogens of M166 and Y95, respectively.

These nitrogen atoms form the “oxyanion hole,” stabilizing the tetrahedral intermediate during the reaction. Notably, fluctuations in these distances are larger than those observed for the similar tetrahedral intermediate in Figure 2B.

Another significant event is the transfer of the excess proton from the water molecule to H242 at approximately 70 ps, resulting in a protonated H242 (N_*ϵ*_).

Figure 4C shows free energy changes and uncertainty estimation during the simulation. The red dots represent the average free energy difference relative to the initial state, with error bars indicating the estimated standard deviations. The free energy barrier for this step ranges from 20 kcal/mol (lower bound path) to 41 kcal/mol (upper bound path).

During this simulation step, only the distance between the water oxygen and the carbonyl carbon in the covalent intermediate (MHET C-Water O) is constrained using our enhanced sampling technique (Section QuantaMind MD Simulation), with no additional biases applied. In the final step, cleavage of the ester bond in the AEI intermediate (Figure 5A) produces terephthalic acid and the original catalytic triad. This transition is accompanied by increased fluctuations in the M166 and Y95 hydrogen bonds (Figure 5B), indicating weaker hydrogen bonds between the product and the “oxyanion hole.” A notable event is the proton transfer from HIP242 to S165, completing restoration of the original catalytic triad.

Figure 5C shows free energy changes and uncertainty estimation during the simulation. The red dots represent the average free energy difference relative to the initial state, with error bars indicating the estimated standard deviations. The free energy barrier for this step ranges from 5 kcal/mol (lower bound path) to 17 kcal/mol (upper bound path).

During this simulation step, only the distance of the ester bond in the covalent intermediate (MHET C-S165 O) is constrained using our enhanced sampling technique (Section QuantaMind MD Simulation), with no additional biases applied.

## Discussion and Future Work

In this study, we employed QuantaMind MD, a MLFF trained on diverse high-level quantum mechanical (QM) data with an emphasis on transition-state regimes, to simulate the complete catalytic cycle of the PETase enzyme over 400 ps of simulation time. Our simulations capture key proton transfer events throughout the reaction, consistent with mechanisms reported in previous studies.^19–26^ Distance analysis underscores the critical role of the backbone amide nitrogens of Y95 and M166, which function as the “oxyanion hole” by forming hydrogen bonds that stabilize the carbonyl oxygen atom throughout the reaction.

The free energy analysis provides quantitative insights into the reaction activation energies. Our simulations estimate free energy barriers of 22 and 20 kcal/mol for the acylation and deacylation steps, respectively, following the lower bound path. These values are relatively close to the estimated experimental value of 18.7 kcal/mol^2,24^ and the calculated value of 21.1 kcal/mol by Boneta et al.^24^ Nevertheless, we acknowledge several limitations affecting the accuracy of the free energy barrier estimation. The large error bars primarily stem from the relatively short simulation time, necessitating longer simulations for QuantaMind to sample sufficient configurations and converge the mean force. Additionally, in the acylation step, the nucleophilic attack and C-O bond cleavage may proceed in a concerted rather than sequential fashion.^19^ Thus, a two-dimensional distance scan is required to construct the full potential energy surface and determine the lowest energy path. A similar approach applies to the deacylation reaction’s two sub-steps.

Finally, as a DL-based model, QuantaMind relies heavily on its training data. While the training of QuantaMind significantly considers transition-state geometries, ^34^ additional training data would be beneficial, particularly for estimating energy barriers. The transition-state structures sampled in this work provide a valuable source of geometries that can be further optimized at the DFT level. These calculations can serve as additional training data to iteratively enhance the QuantaMind model, improving its accuracy and applicability to serine hydrolase-related biomolecular systems.

Overall, this work demonstrates the versatility of QuantaMind MD as a powerful tool for simulating enzymatic reactions in explicit water solvents with ab initio accuracy. Beyond PETase, this framework can be readily extended to a wide range of enzyme classes. Looking forward, we envision that this MLFF-based methodology will not only complement but ultimately supersede traditional QM/MM approaches, eliminating the need for artificial system partitioning while retaining quantum-level accuracy at accessible computational cost. With continued refinement, QuantaMind MD has the potential to establish a new paradigm for studying enzymatic catalysis and biomolecular reactivity.

## Materials and Methods

### Simulation System Preparation

We began with the crystal structure of the leaf-branch compost cutinase (LCC) variant (PDB ID: 7W44),^13^ retaining only the protein structure after removing small molecules and solvents. To incorporate the MHET molecule, we used the crystal structure from PDB ID: 7VVE, which includes MHET and a mutant S165A. Specifically, we aligned the backbone atoms of the catalytic triad from PDB 7VVE to PDB 7W44 and applied this transformation matrix to the MHET molecule in 7VVE. The transformed MHET molecule was then processed with OpenBabel^35^ to add hydrogen atoms.

We utilized AmberTools^36^ for further processing of the protein and MHET structure. Initially, we used the pdb4amber tool to clean the PDB file and add any missing hydrogen atoms. Subsequently, tLEAP was employed to combine the protein PDB and the MHET molecule, followed by solvation with TIP3P water using a 5 Å buffer distance.

### QuantaMind MD Simulation

Following system preparation, we optimized the structure using QuantaMind with the Limited-memory Broyden-Fletcher-Goldfarb-Shanno (L-BFGS) optimizer, performing 1,000 optimization steps.^37,38^ The optimized system was then simulated with QuantaMind under periodic boundary conditions within the NVT ensemble, employing a 0.5 fs time step at 300 K.

To enhance the simulation process, we introduced a biased potential in addition to the potential energy surface predicted by QuantaMind. The biased potential is defined as follows:

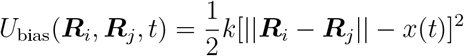

Here, *i* and *j* represent the indices of the atom pairs of interest in the system. The atom pairs are dependent on the specific reaction simulated, as elaborated below. ***R***_*i*_ and ***R***_*j*_ are the 3D coordinates of the respective atoms. The constant *k* = 10.0 eV Å^*−*2^ is a strong force constant, ensuring the distance ||***R***_*i*_ − ***R***_*j*_|| throughout the simulation. The target distance *x*(*t*) evolves according to:

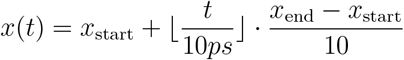

Here *x*_start_ is the initial distance between atoms *i* and *j*, and *x*_end_ is the desired final distance. The values *i, j* and *x*_end_ are determined based on prior chemical knowledge, detailed as follows:

*Acylation reaction-nucleophilic attack: i* and *j* are indices for the oxygen atom of S165 and the carbonyl carbon of the reactant, respectively, with *x*_end_ = 1.45Å.

*Acylation reaction-bond breaking: i* and *j* are indices for the oxygen atom of the partially formed ethylene glycol and the carbonyl carbon of the reactant, respectively, with *x*_end_ = 3.0Å.

*Deacylation reaction-nucleophilic attack: i* and *j* are indices for the oxygen atom of the water reactant and the carbonyl carbon of the reactant, respectively, with *x*_end_ = 1.45Å.

*Deacylation reaction-bond breaking: i* and *j* are indices for the oxygen atom of S165 and the carbonyl carbon of the reactant, respectively, with *x*_end_ = 3.5Å.

### Free Energy Barriers Estimation

Inspired by thermodynamic integration,^39^ we estimate free energy changes using the equation:

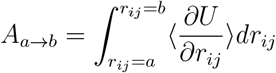

Here, *A*_*a→b*_ represents the free energy change as the distance between two atoms of interest, *r*_*ij*_ transitions from *a* to *b*. The term 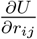is approximated by:

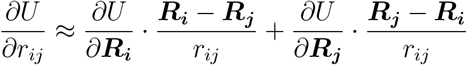

This approximation involves the sum of the projections of the unbiased forces on atoms *i* and *j* onto the vector ***R***_***i***_ − ***R***_***j***_, where ***R***_*i*_ and ***R***_*j*_ are the 3D coordinates of the respective atoms. Using multiple frames obtained from QuantaMind MD simulations, we calculate averages 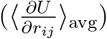 and standard deviations 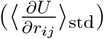, approximating the distribution of 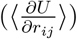with a Gaussian distribution:

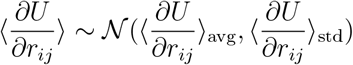

The Gaussian distributions at each *r*_*ij*_ are then propagated through numerical integration to estimate the distribution of the free energy change *A*_*a→b*_.

## Data Availability

The source code used to perform QuantaMind MD simulation is available at https://github.com/MoleculeMindOpenSource/QuantaMind. The molecular dynamics trajectories generated in this work are accessible via Zenodo at https://zenodo.org/records/17217376.

